# A mechanistic model for the evolution of multicellularity

**DOI:** 10.1101/115832

**Authors:** André Amado, Carlos Batista, Paulo R. A. Campos

## Abstract

Through a mechanistic approach we investigate the formation of aggregates of variable sizes, accounting mechanisms of aggregation, dissociation, death and reproduction. In our model, cells can produce two metabolites, but the simultaneous production of both metabolites is costly in terms of fitness. Thus, the formation of larger groups can favor the aggregates to evolve to a configuration where division of labor arises. It is assumed that the states of the cells in a group are those that maximizes organismal fitness. In the model it is considered that the groups can grow linearly, forming a chain, or compactly keeping a roughly spherical shape. Starting from a population consisting of single-celled organisms, we observe the formation of groups with variable sizes and usually much larger than two-cell aggregates. Natural selection can favor the formation of large groups, which allows the system to achieve new and larger fitness maxima.

## 1 Introduction

The formation of new and higher levels of biological organization is a common place in evolutionary biology (Smith and Szathmáry 1997; Szathmáry 2015), being one of the most important transitions the emergence of multicellularity from single-celled organisms. Nowadays, it is well documented that multicellularity has independently occurred in different clades and sometimes even within the same clade (Grosberg and Strathmann 2007; Bell and Mooers 1997; Niklas and Newman 2013). The understanding of the mechanisms supporting that process and, in a broader context, the increase of complexity is a matter of longstanding debate in the scientific community (Smith and Szathmáry 1997; Bonner 1998).

The most prominent feature of these events refers to the transition of independent replicators to associate and form a higher level unit (Smith and Szathmáry 1997). As organismal complexity increases so does the potential to develop division of labor among its lower-level units, which allows the higher-level units to be more efficient under specific conditions, resulting in a fitness advantage. Complexity increases with size as a result of a greater degree of cooperative division of labor within larger entities (Bell and Mooers 1997), a feature also observed on economic grounds (Ghiselin 1978). But this also holds at the framework of competitive interactions among constituents of the system. Indeed, this has been observed in ecological communities, in which larger areas can harbor more species (Rosenzweig 1995; Turner and Tjørve 2005; Campos et al. 2013; Campos et al. 2012).

As aforementioned, multicellularity have evolved in many distinct occasions and through different mechanisms and modes of development, making use of different aspects of cellular biology (Niklas and Newman 2013). In order to multicellular form evolve, there are some requisites that must be followed. For instance, there must exist a fitness transfer from a lower-level biological organization (single cell) to a higher-level biological organization (group) (Wolpert and Szathmáry 2002; Folse III et al. 2010; Michod 2007). This is achieved if, besides cell-to-cell adhesion, some form of communication arises, e.g. biochemical signalling, allowing the cells to communicate and cooperate (Niklas and Newman 2013). A natural and following stage of the process is cell specialization (division of labor), which can minimize any possible conflict among competing physiological processes (Ispolatov et al. 2012). Though it is important to highlight that there are benefits at the earlier stages of the emergence of multicellularity even with undifferentiated cells, as increased group size can enhance predation avoidance (Boraas et al. 1998; Grosberg and Strathmann 2007), or even create a buffered environment within a group, which can favor the sharing of secreted public goods (Koschwanez et al. 2011). Thus, undifferentiated multicellularity amounts to an additional advantage for the evolution of specialized cells (Willensdorfer 2008). Moreover, the insight that the size of an organism affects its ability to evolve new specialized cells is vital for our understanding of how complex multicellular life evolved. The size of a multicellular organism affects its fitness landscape. This augment entails an increase in the dimensionality of phenotype space, and the generation of a fitness landscape with new fitness maxima ensues (Willensdorfer 2008; Ispolatov et al. 2012).

Here we explore the conditions in which the formation of groups of arbitrary sizes is favored. We adopt a mechanistic approach to investigate the evolution of multicellularity. This approach takes into account quite distinct mechanisms driving the dynamics of group size, such as aggregation, dissociation, reproduction and cell death. In the model, there is an important interplay between natural selection and dynamics of group growth as the size of groups strongly impact rates at which cells divide. In the current work, our aim is to observe the emergence of aggregates of arbitrary size thus allowing us to understand the underlying mechanisms that can limit the size of evolving organisms. The problem is simulated within the framework of kinetic theory, where finite populations are assumed and thereby stochastic components of the evolutionary dynamics are naturally incorporated. The problem is formulated within the context of existing tradeoffs. The cells are genetically identical and can complete two tasks, for instance, the production of two metabolites. These two tasks are poorly compatible, which makes room for the appearance of division of labor as bigger organisms arise. The time scale at which signalling processes among cells in a group occur is much faster than the mechanisms directly affecting group size dynamics, and so one important premise of the modelling is that internal arrangement of cells in a groups is the one that maximizes organismal fitness. Note that the establishment of a reproductively integrated phenotype is one of the requirements for multicellularity (Niklas 2014). This coordination among group members is seen in cellular evolution (Dekel and Alon 2005) as well as in social evolution (Duarte et al. 2012). Once a multicellular organism has been formed, that allows for selection of further size increase, which in its turn will require an increase in complexity, i.e., a further development of division labor through cell specialization. This is dubbed as the size-complexity rule (Bonner and Brainerd 2004). Of course, each newly assembled cell does not add up a new different function. In spite of keeping the number of functions fixed and equal to two, here we show that there exists a selection for “intermediate” organismal size, as a change in organismal size affects the organism phenotypic space, thus allowing the search for new local maxima.

The paper is organized as follows. In Section II the model is described. Section III presents the theoretical development as well as discusses in detail the formulation which allows us to simulate the problem as a set of chemical reactions, and the type of structures considered in the current work. Section IV shows the simulation results. And finally, in Section V we present our concluding remarks.

## 2 The Model

We propose a model of cell aggregation in which two different scenarios for the aggregate shape are addressed. The population is finite and consists of cells that can reproduce, die and lump together giving rise to aggregates of different sizes. The cells can also exist in its unicellular form. The handling of the formation of aggregates is a quite difficult matter as these aggregates can produce quite distinct and complex structures. At this point some simplification is needed. In a recent study, Ispolatov et al. formulated a model for formation of aggregates of up to two cells (Ispolatov et al. 2012), i.e. cells could either form two-cell aggregates or remain in its unicellular form. The generalization of such analysis to aggregates of arbitrary sizes brings about some difficulties. The first concerns the assumptions underlying fitness optimization within the groups. Instead of maximizing cell’s fitness, as originally supposed, we consider that the organismal fitness is the quantity to be optimized or likewise the per capita reproductive rate. The second and troublesome issue is that by designing organisms comprised of an arbitrary number of cells the group structure can no longer be disregarded. Though, the assessment of groups of arbitrary structures is prohibitively complex. Even in the case of a single mechanism, such as aggregation, the problem is already quite complicated (Krapivsky et al. 2010). For the sake of feasibility, the current analysis restrains the study to the formation of multicellular organisms that assume either a spherically symmetric structure or a linear chain structure.

The cells are endowed with the capability to perform two tasks. Let us think about of the production of two metabolites, hereafter denoted by *X* and *Y*. The production of these metabolites brings about a benefit *B*(*x*, *y*) and a cost *C*(*x*, *y*). The production of both metabolites can not grow without a bound, due to the existence of a tradeoff. As such, the production of both metabolites in considerable amounts inflicts a huge cost to the cell. However, both metabolites are considered to be essential for the cells. With this aim, the benefit *B*(*x*, *y*) and the cost *C*(*x*, *y*) are written as

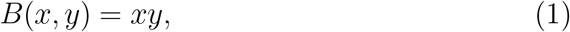

and

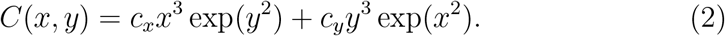

The form of Eqs. (1) and (2) ensures that the cost *C*(*x*, *y*) grows faster than the benefit *B*(*x*, *y*) in any direction of change in *X* and *Y* (Ispolatov et al. 2012). The prominent increase of the cost *C*(*x*, *y*) with the expression level of a trait reflects the fact that the cell has limited resource. For instance, the production of a given protein results in reduced capability to produce other proteins, as claimed by Dekel and Alon (Dekel and Alon 2005). The latter study also observes that cells rapidly evolve in order to reach optimal expression levels, thus maximizing the cost-benefit problem.

A single cell produces both metabolites *X* and *Y*, which, in their turn, are consumed by the cell itself. When forming aggregates, the metabolites are equally shared but the cost of metabolite production is individual. Therefore, the per capita reproduction rate, equated as fitness, in an *n*-cell aggregate is

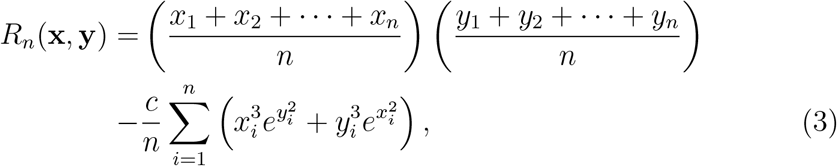

where *x*_*i*_ is the amount of metabolite *X* produced by the cell *i*, while the vector **x** stands for the set of variables (*x*_1_, *x*_2_, …, *x*_*n*_), analogous definitions holding for *y_i_* and y. For simplicity, we have considered *c*_*x*_ = *c*_*y*_ = *c*. A key assumption of the modeling is that, for any aggregate size, the metabolism of the cells in the aggregate always works at optimized levels, i.e., at level 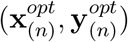 that optimizes the benefit-cost function as defined in Eq. (3), where the subscript *n* has been used because the optimal level of metabolites may depend on the aggregate size, as we shall see in the sequel.

The model also assumes that the strength of the linkage between cells is determined by the stickiness *σ*, 0 = *σ* = 1.

## 3 Theoretical development

For unicellular organisms, the reproduction rate *R*_1_ reduces to

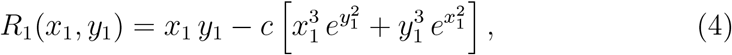

being optimized at level 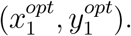 Due to the symmetry of the above function, 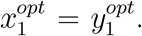 Denoting this optimal value by *χ*_1_, the reproduction rate of the single-cell can be maximized by deriving the function *R*_1_(*χ*_1_, *χ*_1_) with respect to *χ*_1_ and equating the result to zero, which yields a transcendental equation. For instance, if *c* = 0.04 we have 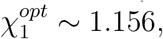 so that 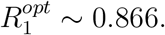

For two-cell aggregates, we expect division of labor to show up, as it is costly to keep the simultaneous production of the two metabolites. Through the assumption of complete symmetry between the two cells, the maximum fitness should be attained when 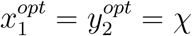 and 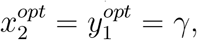 so that 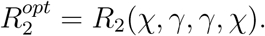 For instance, when *c* = 0.04, the optimization occurs at *χ* ∼ 4.16 and *γ* ∼ 0.00, yielding 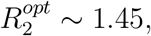 as shown in Ref. (Ispolatov et al. 2012) and endorsed by our results. Obviously, a complementary configuration *χ* ∼ 0.00 and *γ* ∼ 4.16 also yields the same fitness, 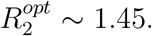 This result supports the relevance of division of labor, inasmuch as the optimal case is obtained when cell 1 produces just the metabolite *X* while cell 2 produces just the metabolite *Y*, or the converse. Moreover, note that the reproduction rate per cell in the two-cell aggregate is larger than in the single-cell state. In this case, natural selection favors the formation of the two-cell aggregate in comparison to the single-cell state, endorsing the evolutive advantage of labor division and, eventually, cell specialization.

We should note that the maximization of the per capita reproduction rate in a *n*-cell aggregate, *R*_*n*_, is the same as the maximization of the organismal fitness, which is just *nR*_*n*_. The maximization of fitness of the whole organism is not only made for practical reasons but also because, as part of a higher level biological organization, the target of selection is no longer the cell itself but the entire group (Niklas 2014; Willensdorfer 2008). By extending the study to larger aggregates we are faced with two distinct situations depending on the parity of the number of cells in the aggregate, *n*. When *n* is even, one can grasp from the previous considerations about symmetry in the two cell aggregate that the optimal configuration corresponds to half of the cells producing the metabolite *X* and essentially performing no production of *Y*, while the remaining ones do the opposite, namely just produces *Y*. For instance, we can say that the optimal fitness is obtained when

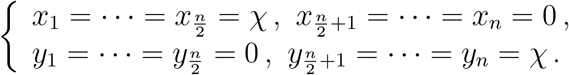

Under this configuration, the per capita reproduction rate becomes

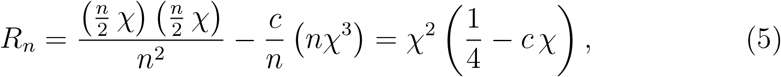

which is optimized at *χ* = (6*c*)^−1^, yielding 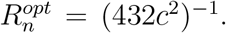 Thus, for *n* even, *R*_*n*_ becomes independent of *n*.

For an odd number of cells the situation is more subtle. At a first glance, a natural choice is the situation in which 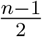 cells produce metabolite *X* and no metabolite *Y*, 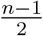 cells produce metabolite *Y* and no metabolite *X* while the remaining cell produces an equal amount of both metabolites, namely:

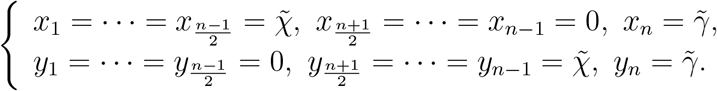

Indeed, for high values of the cost parameter *c*, the above configuration is the optimal one. Nevertheless, in the range considered here, 0 < *c* < 0.1, this can be the optimal configuration only for *n* = 3 and *n* = 5 and, even though, restricted to a small range of *c*. Yet, for *n* ≥ 7 the optimal fitness is reached when 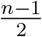 cells produce one of the metabolites, and the 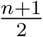 remaining cells produce the other metabolite, i.e.,

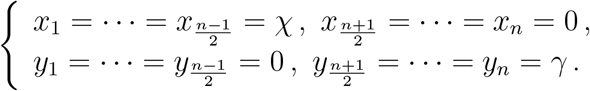

In such a case the reproductive rate becomes:

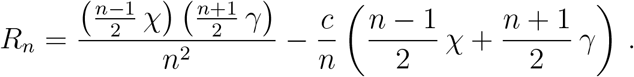

Maximizing the latter with respect to the parameters *χ* and *γ* we end up with the following values:

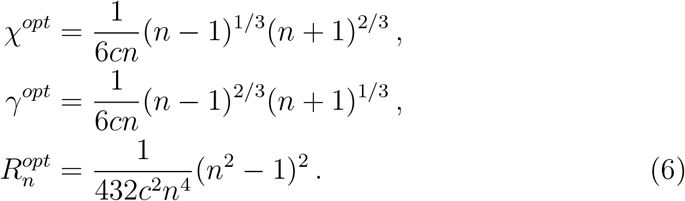

Note that *χ^opt^* > *γ^opt^*, which should be expected due to the assumption that in the aggregate there are more cells producing *Y* than cells producing *X*. Although the rate (6) is not the optimal one for the special cases *n* = 3 and *n* = 5 in the full range 0 < *c* < 0.1, on average the difference is not significant. Moreover, we have checked that no qualitative behaviour is compromised by the interchange of both rates. Therefore, in the sequel, we shall use the fitness presented in Eq. (6) whenever *n* is odd and *n* ≥ 3.

### 3.1 Stochastic Dynamics

The problem here addressed is studied through extensive computer simulations. The model dynamics can be written as a set of chemical reactions that considers all the possible state transitions, whose rates are determined by the population densities of the aggregates and reaction constants. The dynamics includes the following possible events: (i) aggregation; (ii) dissociation; (iii) cell death (apoptosis) and (iv) cell reproduction. To simulate the model dynamics we employ the Gillespie Algorithm (Gillespie 1976). According to the method, time between events (a chemical reaction) is exponentially distributed, and the likelihood that a given reaction occurs is just its rate divided by the total rates of all possible reactions.

Below we present and enumerate the set of reactions that represents the possible transitions in the model. Hereafter *A*_*i*_ denote an aggregate of size *i*, and the quantities k’s refer to the reactions constants. Due to the specificities of the two structures here investigated, the set of possible reactions and the estimate of their rates will be presented separately. Though in both types of structure these reactions include the processes of aggregation, dissociation, cell death and reproduction.

#### 3.1.1 The linear model

In the linear model, the group of cells forms a chain. The set of allowed reactions and estimate of their rates are described below and sketched in Fig. 1.

i. Aggregation:

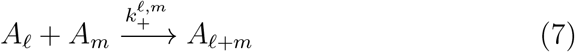

An *ℓ*-cell aggregate can combine with an *m*-cell aggregate to form an aggregate of size *ℓ* + *m*. Due to the linearity of the chain the binding force between the two chains is proportional to the square of the stickiness *σ*. Therefore,

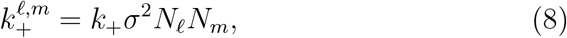

where *k*_+_ is the rate of aggregation, and *N*_*k*_ stands for the number of *k*-cell aggregates.
ii. Dissociation:

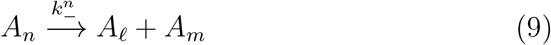

An aggregate of size *n* can give rise to two aggregates of sizes *ℓ* and *m*, with the constraint *n* = *ℓ* + *m*. The rate of the aforemention process is determined by

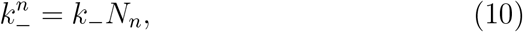

where *k*_−_ is the rate of dissociation of the aggregate. For the linear structure the dissociation of every pair of cells in the aggregate is equally likely, which means that *ℓ* can take values *ℓ* = 1, …, *n* − 1. Note that the dissociation does not depend on the stickiness as also assumed in Ref. (Ispolatov et al. 2012).
iii. Cell death. The process of death is depicted as

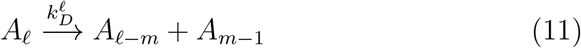

where *m* is the position of cell to be removed, i.e. *m* = 1, …, *ℓ*. If the cell to be eliminated is located at the edges then one of the two terms in the right of the reaction will equal 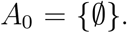 In this case only one chain of size *ℓ* − 1 will result from the process. One cell in an *ℓ*-cell aggregate dies at rate

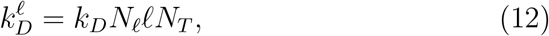

where *k*_*D*_ is the rate of death per cell, and NT means the total number of cells in the population and displays an important regulatory role for the system size.
iv. Cell reproduction:

– With probability 1 − *p*_*single*_ the daughter cell remains attached to the aggregate

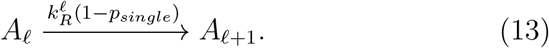
–With probability *p*_*single*_ the daughter cell detaches from the aggregate

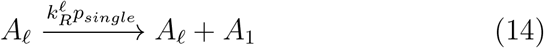

In case a cell reproduction happens, one has two possibilities: either the cell remains in the aggregate, thus increasing its size by one, or the daughter leaves the chain, thus forming an one-cell aggregate. The former event occurs at rate 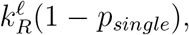 whereas the second event takes place at rate 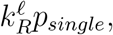 where *p*_*single*_ denotes the probability to leave the aggregate. In the simulations we took *p*_*single*_ = 1 – *σ*^2^, since an increase in the stickiness leads to a decrease in the probability of the new cell leaving the group. The rate 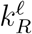 is given by

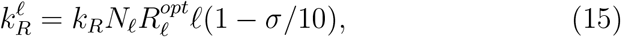

*k*_*R*_ stands for the reproduction rate, 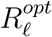 is the fitness per capita of the aggregate with *ℓ* cells, and the factor (1 − *σ*/10) represents the cell’s cost due to stickiness.

**Figure 1:**
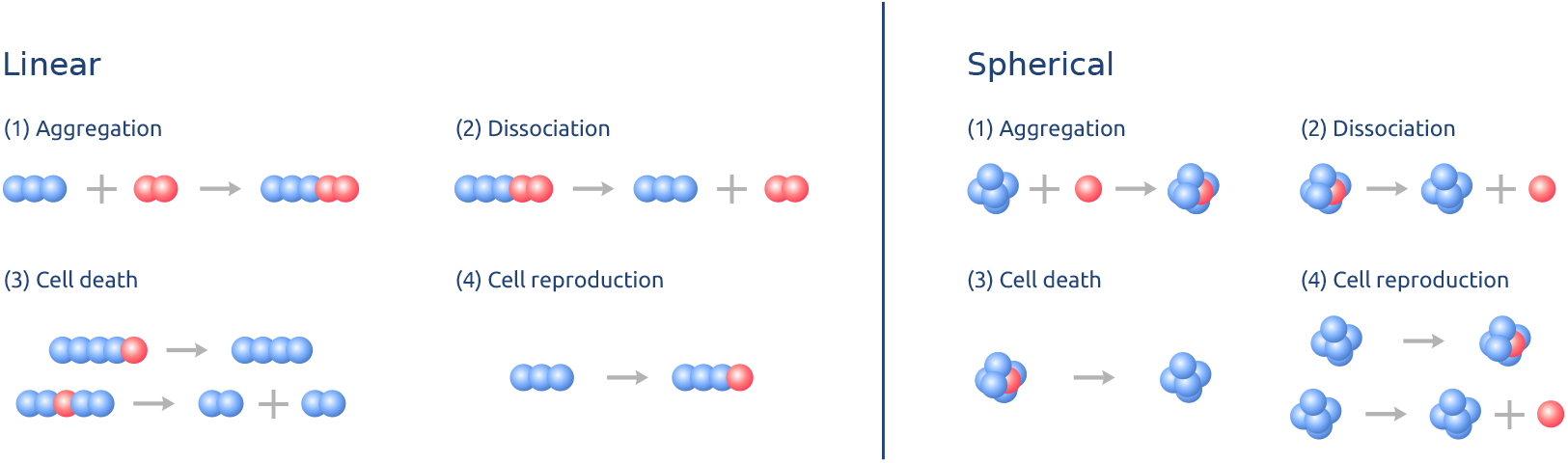
Sketch representing the several processes included in the model.

The above set of reactions (transitions between states) is performed through the use of the Gillespie Algorithm.

#### 3.1.2 Compact aggregates – spherical symmetry

Here we consider two distinct shapes for the aggregates. In the case just considered the aggregate forms a linear chain. Now, we will assume that the cells form a very compact aggregate, which can be coarsely-grained portrayed as groups of cells exhibiting a spherical symmetry. The rates of some of the allowed reactions are calculated differently in both geometries, as shown in the sequel and depicted in Fig. 1.

i. Aggregation:

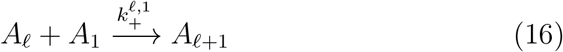

For the spherical structure, an *ℓ*-cell aggregate can only combine with an one-cell aggregate to form an aggregate of size *ℓ* + 1. It is assumed that the event of aggregation involving two aggregates of large size is very unlikely, as it would be very costly for the aggregate to rearrange itself in a spherical shape. The likelihood that an one-cell aggregate collides with an *ℓ*-cell aggregate is proportional not to the number of cells in the group but rather scales with *ℓ*^2/3^ (surface area). In this case, the reaction rate for the above process is calculated as

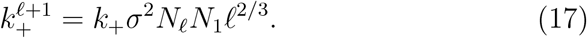

In the linear model it is implicit a constant reaction kernel, i.e., the reaction rates are the same regardless the size of the aggregate (Krapivsky et al. 2010), while for the spherical model the reaction rates depend on the mass of the aggregate. Though, it is not a product kernel as it does not scale with the number of cells in the aggregate, but rather with the surface area.
ii. Dissociation:

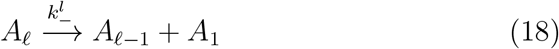

For the spherical model we assume that an aggregate of size *ℓ* can give rise to two aggregates, one of size *ℓ* – 1 and the other of size 1, the rate of this process being

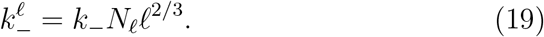

In comparison to the linear model there is an additional factor, *ℓ*^2/3^, since events of dissociation are expected to occur in cells at the surface of the group.
iii. Cell death:

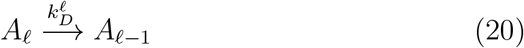

One cell in an *ℓ*-cell aggregate dies giving rise to an *ℓ* – 1-cell aggregate. As the group forms a compact organism it is assumed that events of cell are unlikely to produce a dissolution of the group. This process takes place at rate

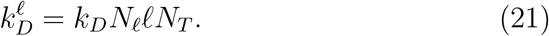
iv. Cell reproduction:

- With probability 1 − *p*_*single*_ the daughter cell remains attached to the aggregate

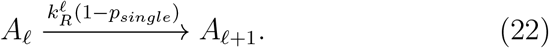
- With probability *p*_*single*_ the daughter cell detaches from the aggregate

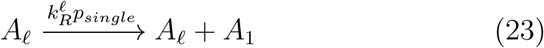

The same situation as the one for the linear model is considered here. The former event occurs at rate 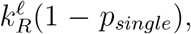 whereas the second event takes place at rate 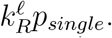 Again, *p*_*single*_ is the probability to leave the aggregate. The rate 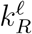 is given by

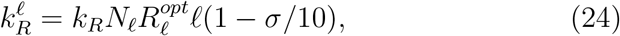

where, again, *k*_*R*_ denotes the reproduction rate, 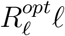 is the total fitness of the aggregate and the factor (1 − *σ*/10) represents the cell cost due to stickiness.

## 4 Simulation Results

Figure 2 shows the average trajectory for the mean group size and the maximum group size with the number of iterates for both linear and spherical models. Each iterate corresponds to one event (reaction). In both models, linear and spherical, one clearly observes the formation of relatively large groups. For this set of par ameter values, groups com prise up to 30 cells at equilibrium stage, though the average group size is consider ably smaller, as groups of single ce lls are abundant due to the mechanism of dissociation and reproduction. One interesting feature is the existence of an overshooting for the linear model, whereas the approach to stationarity in the spherical model is smoother. One basic difference between the linear and the s pherical models is that the former allows the combination of large aggregates, while in the spherical model an aggregate can only change its size by one unit at a time. This may explain the overshoot in group size in the earlier stages of the evolution in the linear model.

**Figure 2:**
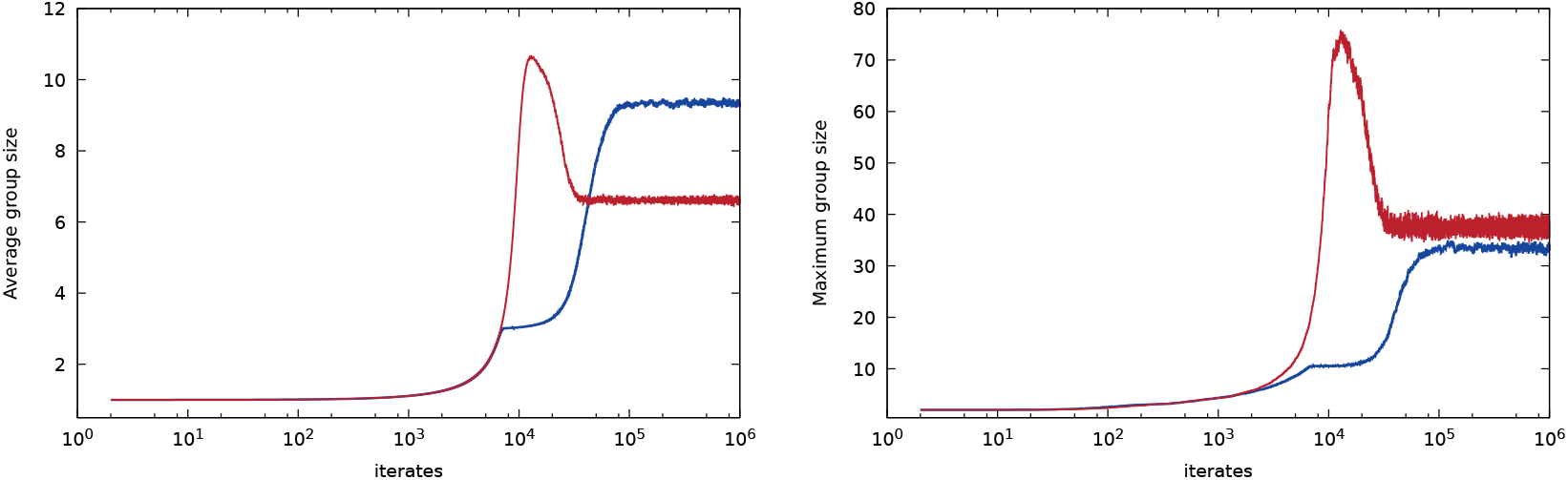
Evolutionary trajector ies for mean group size (left panel) and largest group size (right panel). The red lines represent the results for the linear model whereas the blue lines refer to the s pherical model. The parameter values are *k*_+_ = 0.01, *k*_−_ = 0.01, *k*_*R*_ = 0.01, *k*_*D*_ = 10^−5^, *σ* = 0.5 and *c* = 0.04. The res ults correspo nd to an ave r age over 100 independent runs.

Distributions of group size at stationarity for several values of stickiness are presented in Figure 3. The outcomes for both models are qualitatively similar, and even quantitatively the patterns do not differ much. As expected, the increase of stickiness favors the formation of larger groups despite its increased fitness cost. For large stickiness, the probability distribution of group sizes is nearly flat over a broad range of size and then abruptly drops when the group size passes a threshold, especially in the spherical model. The cut-off size increases with stickiness. For the set of parameters here considered, group sizes can reach values up to few hundred cells. In general, the probability distribution for the linear model is less flat. There are some important features to point out that are corroborated from these results. First it is important to mention that this is not a simple model of aggregation. Indeed, aggregation is only one of the four mechanisms that rule the evolutionary of the organisms, and it is also not the dominant one. From Fig. 3 one observes that the steady-state probability distribution of aggregate size encompasses aggregates of variable size. This is clearly contrary to what is observed in simple models of aggregation, especially those considering irreversible aggregation, where the distribution would converge to a delta function for a finite number of monomers (cells) (Krapivsky et al. 2010). Models of aggregation with input also differ from our approach. In those models, the tail of the aggregate sizes at stationarity is expected to decay as *n*^−3/2^ (Krapivsky et al. 2010), where *n* stands for group size. In Fig.3 the dashed straight line, which serves as an eye-guide, describes a power-law like *n*^−3/2^, and it is clearly distinct from the tail of the distribution generated in our processes, which exhibits a much steeper decay. All these together, evince that the process of augmentation of group size is not only determined by the process of aggregation. Indeed, even the process of input, appearance of new minimal units (cells), is completely ruled by the dynamics of the system itself. At this context, there is an interesting feedback between natural selection and growth dynamics of aggregates, as their size strongly influence the reproductive rate of cells.

**Figure 3:**
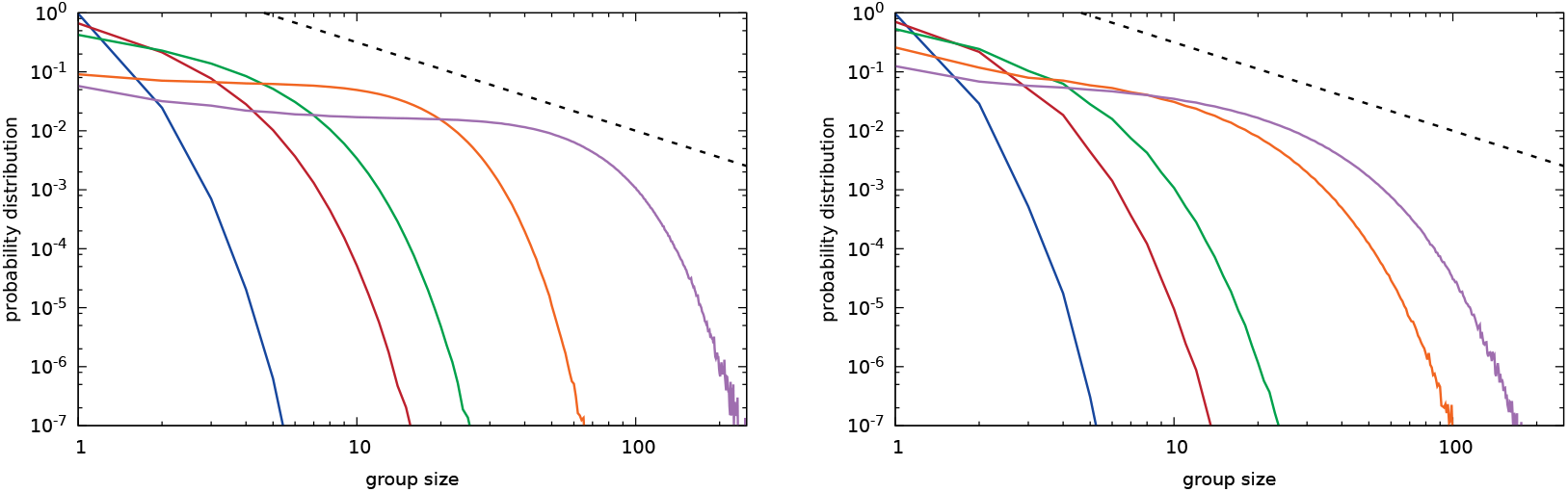
Probability distribution of gro up sizes for the linear (left) and spherical (right) models. The val ues for stickiness are f ro m left to right 0. 01, 0.05, 0.1, 0.5, 0.9. The remaining par ameter values are *k*_+_ = 0.01, *k*_−_ = 0. 01, *k*_*R*_ = 0.01, *k*_*D*_ = 10^−5^, and *c* = 0.04. The results correspond to an average over 100 independent runs. The straight dashed line serves as an eye-guide describing a power-law with exponent −3/2, which is expected to desc ribe the tail of the distribution in a system with only the mechanism of aggregation with in put.

Next, in Figure 4 a heat map of the average and largest group sizes in terms of the cost, *c*, and stickiness, *σ*, is shown. We clearly see that a higher cost of producing metabolites yields smaller size groups. In its turn, higher stickiness favors bigger groups. For low values of stickiness we observe that the cost, *c*, plays a minor role and average group size is roughly constant. Although the average group sizes are not quite distinct in the two models, in the linear model the largest groups can be comprised of a relatively higher number of cells, like one hundred or more cells, as achieved for low cost c and high stickiness *σ*. One important remark is that for low cost *c*, the formation of groups of a given size requires a minimal level of stickiness, *σ*, that is larger for the spherical model.

**Figure 4:**
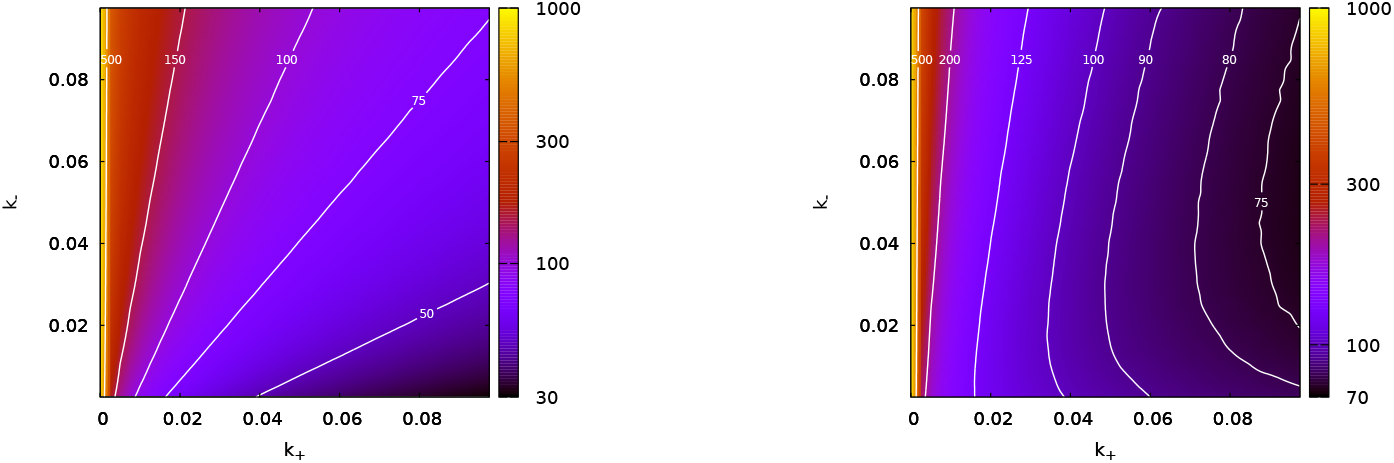
Average group size (upper panels) and maximum group size (lower panels) for the linear (left) and spherical (right) models as a function of *σ* and *c*. The remaining parameter values are *k*_+_ = 0.01, *k*_−_ = 0.01, *k*_*R*_ = 0.01 and *k*_*D*_ = 10^−5^. Each point corresponds to an average over 5 × 10^6^ iterates sampled every 1000 iterates after reaching equilibrium for 16 independent runs.

The dependence of both average and largest group size on the dissociation parameter, *k*_−_, and on the aggregation parameter, *k*_+_, are displayed in Fig. 5. In the linear model (left panel) the variables *k*_+_ and *k*_−_ have opposed effect on both quantities. While group size shrinks as the aggregation rate *k*_+_ decreases, it grows with the reduction of the dissociation rate *k*_−_, i.e., for fixed *k*_+_, the augment of *k*_−_ always results in smaller groups, as we see easily notice from the isoclines. For small values of *k*_+_ a change in the rate of dissociation *k*_−_ has a minor effect. On the other hand, the spherical model exhibits a more counterintuitive and somehow striking behavior. While for small values of the aggregation rate *k*_+_, the augment of the dissociation rate means that smaller groups are formed, for intermediate and large values of *k*_+_ the average group size is peaked at intermediate values of *k*_−_. This feature is highlighted when analyzing the maximum group size for the spherical model, which clearly reveals that the increase of *k*_−_ at large *k*_+_ can produce larger groups. This is particularly interesting as similar effect has been empirically observed in the formation of snowflakes structures in yeast *Saccharomyces cerevisiae* populations (Ratcliff et al. 2012).

**Figure 5:**
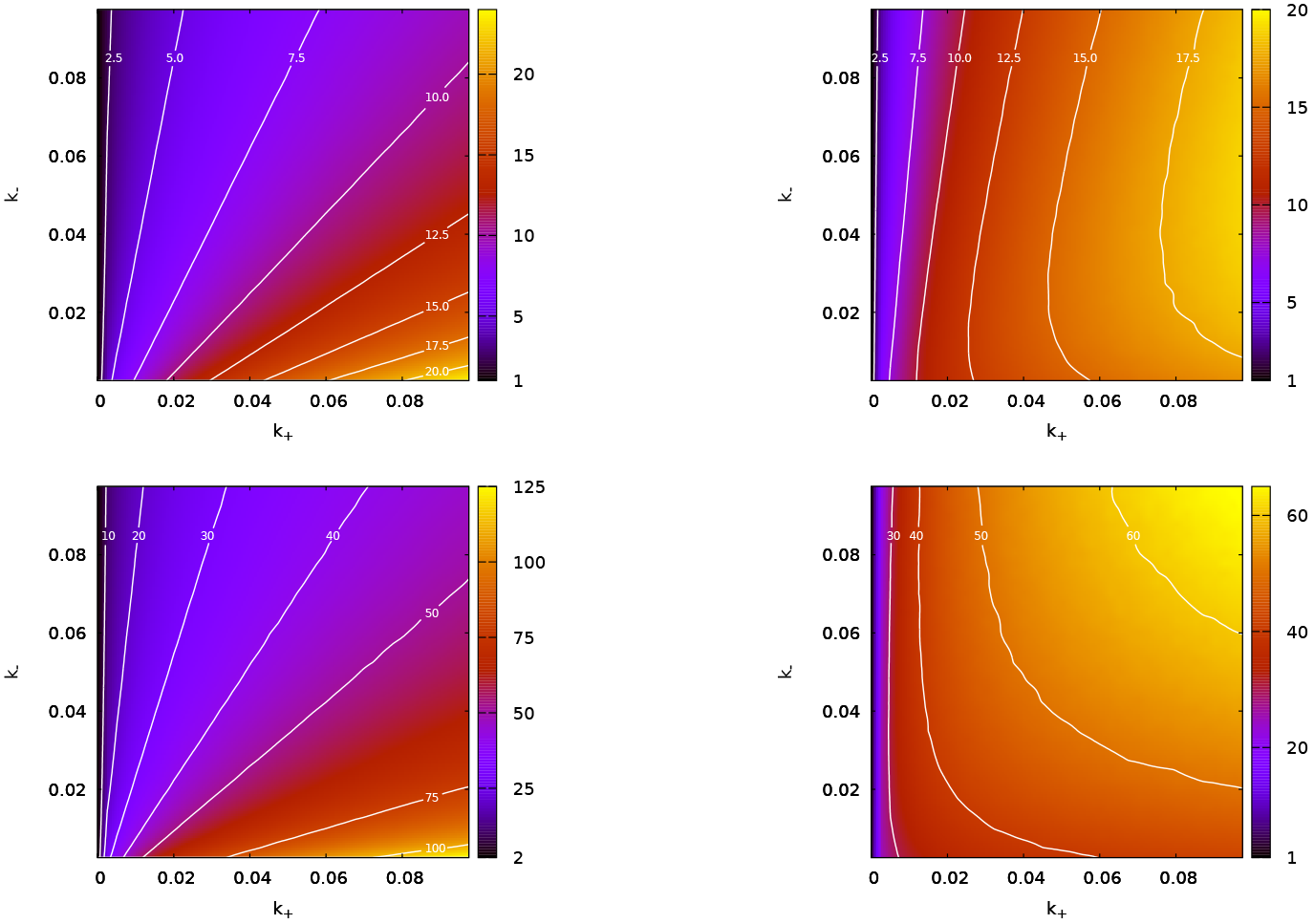
Average group size (upper panels) and maximum group size (lower panels) for the linear (left) and spherical (right) models as a function of *k*_+_ and *k*_−_. The remaining parameter values are *k*_*R*_ = 0.01, *k*_*D*_ = 10^−5^, a = 0.5 and *c* = 0. 04. Each point corresponds to an average over 5 × 10^6^ iterates sampled every 1000 iterates after reaching equilibrium for 16 independent runs.

In the following graphic we show how the number of groups changes in terms of the cost *c* and stickiness *σ*, see Fig. 6. In general, we observe that larger stickiness at fixed cost leads to smaller number of groups. On the other hand, an increased cost *c* has as an outcome that a smaller number of aggregates can be beared by the population at stationarity. A decreased number of groups with increased tradeoff costs is in agreement with previous studies (Amado and Campos 2017; Gavrilets 2010). Except for small costs *c*, the number of aggregates is peaked at small value of stickiness *σ*. Regarding the dependence of the number of groups on the dissociation *k*_−_ and aggregation *k*_+_ rates, the reader is referred to Fig. 7. In the latter we notice that for the linear model there exists a straightforward relation, the number of groups grows with the augmentation of *k*_−_ and the reduction of *k*_+_. Conversely, it dwindles as *k*_−_ is reduced and *k*_+_ is raised. Once again, the spherical model displays a distinguished scenario in comparison to the linear model, as one observes that at intermediate and large *k*_+_ the number of aggregates attains its minimum at intermediate *k*_−_. This is complementary to the outcome presented in Fig. 5, as this region reflects the emergence of larger groups.

**Figure 6:**
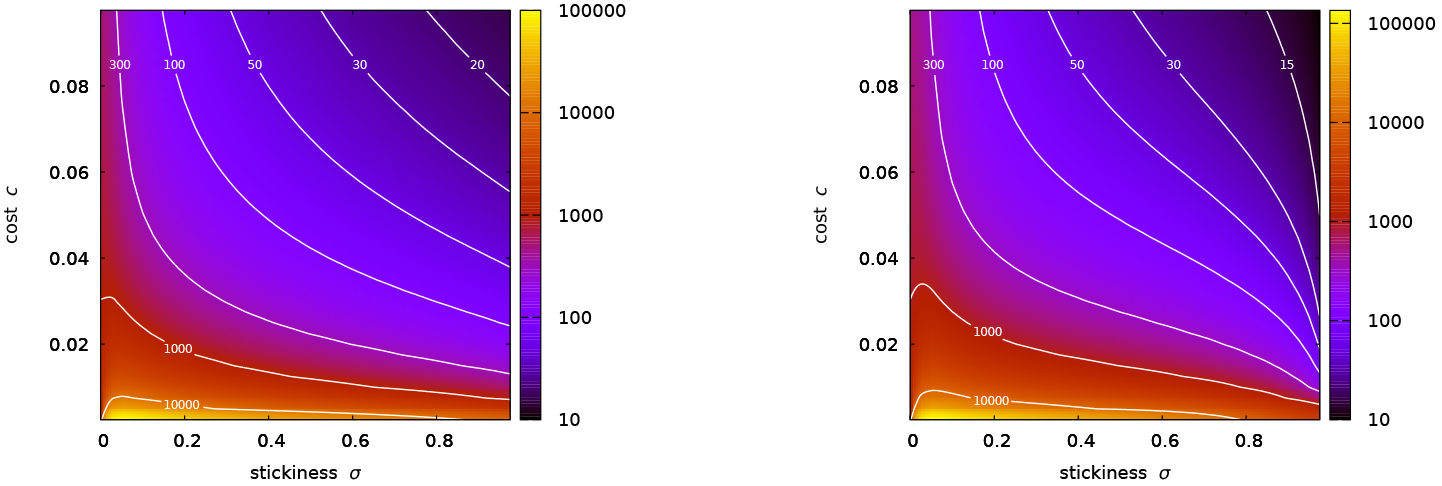
Number of gr oups for the linear (left) and s pherical (right) mode ls as a function of *σ* and *c*. The remaining parameter values are *k*_+_ = 0.01, *k*_−_ = 0.01, *k*_*R*_ = 0.01 and *k*_*D*_ = 10^−5^. Each po int cor responds to an ave r age over 5 × 10^6^ ite r ates sampled every 1000 iterates after reaching equilibrium for 16 independent runs.

**Figure 7:**
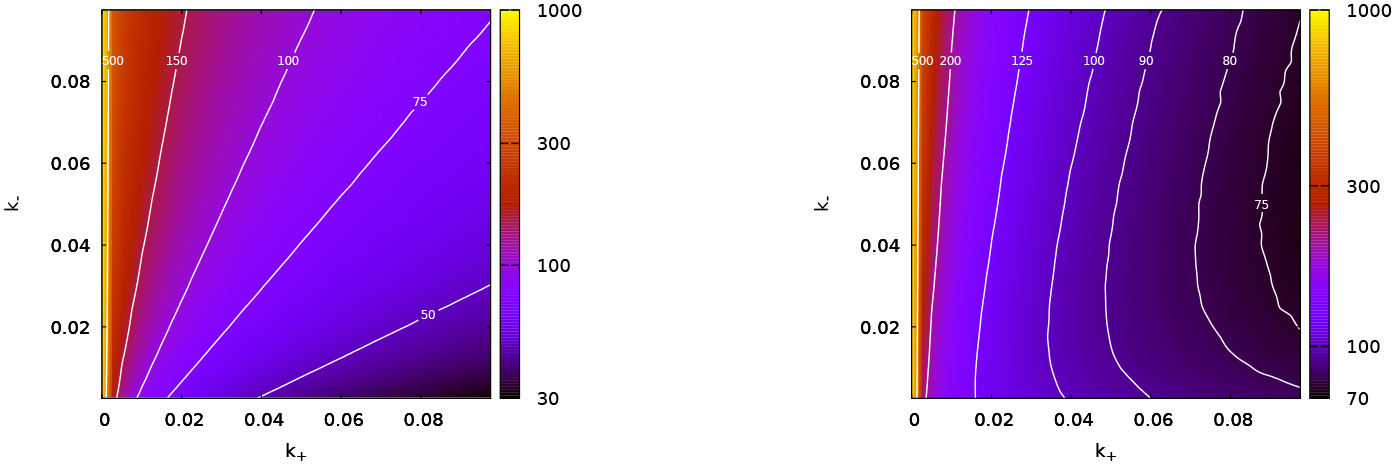
Number of groups for the linear (left) and spherical (right) models as a function of *k*_+_ and *k*_−_. The remaining parameter values are *k*_*R*_ = 0.01 and *k*_*D*_ = 10^−5^, *σ* = 0.5 and *c* = 0.04. Each point corresponds to an average over 5 × 10^6^ iterates sampled every 1000 iterates after reaching equilibrium for 16 independent runs.

## Conclusions

We have investigated how large groups of cells can be formed starting from basic principles such as aggregation, dissociation, cell death and reproduction. Natural selection plays a critical role in the evolution of the system as the group configuration is assumed to rapidly change in order to maximize reproductive rates at the group. In the model, the cells can produce two metabolites but the endeavour of their production is poorly compatible. Though, these metabolites are said to be essential. Also, the production of both metabolites at high amounts can inflict a huge fitness cost to the cell. The appearance of groups can effectively attenuate such incompatibility, as the cells comprising a group can specialize at a given task, thus reducing its fitness cost but at the same time enjoying the benefit of sharing the metabolites produced by other members of the group.

The problem of evolution of complexity, here taken as size of groups, is studied through extensive computer simulations. With this aim, the presentation of the different processes affecting cells and groups (aggregation, dissociation, death and reproduction) are depicted in the form of kinetic chemical reactions, which allows us to make the probability of occurrence (like rate constants) precise. The system is then simulated through the use of the Gillespie algorithm, a powerful Monte Carlo stochastic simulation algorithm to find the trajectory of a dynamical system described by a set of interaction network, such as chemical reactions or even ecological problems (Gillespie 1976). The method is extremely versatile, being of great applicability and not restrictive (Vestergaard and Génois 2015). Starting from a population comprised of single-celled organisms we observe the continuous formation of intermediate and large groups of cells, establishing a stable coexistence of aggregates of varying sizes. As in the modelling groups can be composed of an arbitrary number of cells, it makes room to the appearance of an infinitely large number of possible distinct morphologies, being necessary to settle a choice for the emerging structure of the groups. For the sake of feasibility, here this is addressed by either assuming that groups form linear chains (linear model) or are extremely compact in such way they take a spherical shape (spherical model). In the former case it is assumed that the processes of aggregation, dissociation and cell death enable the group sizes to change discontinuously as these events can either promote linkage of chains of different sizes or promote chain break in different locations of the chain. On the other hand, in the spherical model the transitions are smoother, as all these processes lead to variation in group size not larger than one cell.

In spite of the underlying difference between group shapes, their qualitative behaviors are not quite distinct in respect to the size of the largest groups produced as well as the composition of the population at stationarity. Also, with respect to stickiness and tradeoff cost, in both models we observe that a higher cost will result in smaller groups, and, as expected, the increase of stickiness favors the formation of larger aggregates. Although the cells are assumed to produce only two metabolites, we still observe the formation of aggregates with the number of cells greater than two, meaning an evolutionary advantage towards bigger groups. For two-cell aggregates, there is a clear advantage for the configuration at which one cell produces one metabolite while the other cell produces the second metabolite. For aggregates larger than two, the group will still produce the two metabolites but group expansion reshapes the fitness landscape allowing the system to achieve new and larger fitness maxima. As we know from the study of multicellularity, the introduction of a new cell to a group does not mean that a new function is developed by the organism (Rueffler et al. 2012; Bonner and Brainerd 2004), a fact that also holds in social organisms (Anderson and McShea 2001).

A remarkable difference between the linear and spherical models is found when we analyze how group sizes evolve in terms of the aggregation and dissociation rates. For the linear model, the relation between group size and the rates of aggregation *k*_+_ and dissociation *k*_−_ is straightforward: while the increase of *k*_+_ favors increased group sizes, the augment of *k*_−_ favors smaller groups. Still, for the spherical model, the dependence of groups sizes on both *k*_+_ and *k*_−_ is more subtle. At intermediate and large values of the aggregation rate, *k*_+_, increased values of *k*_−_ can select for larger groups. In a recent paper, Ratcliff et al. found that, after some time of evolution, snowflakes structures developed division of labor with some cells undertaking programmed cell death (apoptosis) (Ratcliff et al. 2012). It is claimed that programmed cell death is evolutionary advantageous since it allows the snowflakes yeast to increase its fecundity. Indeed, they demonstrated that larger average group size and larger rates of apoptosis have coevolved (Ratcliff et al. 2012). In our model, dissociation plays the same role as cell death from the perspective of the group. Therefore, the spherical model, which allows smoother transitions in terms of variation in group size, seems to capture this feature of the empirical observation of the snowflakes yeast experiment (Ratcliff et al. 2012).

Our simulations show that group size evolves until it levels off and the mechanisms that favor group growth like aggregation and reproduction are balanced by the mechanisms supporting disruption. Particularly, it is worth mentioning that as time evolves in the linear chain model the average group size first passes through a peak and then drops to the stationary value. As aforementioned, aggregate sizes can grow further than two cells as an outcome of fitness reshaping which, in its turn, allows the whole organism to explore and find higher levels of fitness, i.e., new arrangements are found in which the process of division of labor can turn even more effective. One may also conjecture that the maximum sizes reached in this dynamics reflects an upper bound for aggregate sizes, and a further growth in the number of cells comprising the organisms requires additional elements, such as the production of a new metabolite or any public good that can be shared and is beneficial to the group, or even some strategy enabling transfer of fitness from the individual level to the group level. In brief, from this point on, more division of labor will be required.

## Acknowledgments

PRAC is partially supported by Conselho Nacional de Desenvolvimento Científico e Tecnológico (CNPq), and also acknowledges financial support from Fundação de Amparo à Ciěncia e Tecnologia do Estado de Pernambuco (FACEPE) under Project No. APQ-0464-1.05/15 and Universidade Federal de Pernambuco Edital Qualis A 2016. CB is thankful to Conselho Nacional de Desenvolvimento Científico e Tecnológico (CNPq) for the partial support. AA has a fellowship from Conselho Nacional de Desenvolvimento Científico e Tecnológico (CNPq).

